# Temporal regulation of motor behavior on a modified forelimb dexterity test in mice

**DOI:** 10.1101/2020.10.18.344507

**Authors:** Hisham Mohammed, Yue Li, Paola Di Grazia, Amanda Bernstein, Sydney Agger, Edmund Hollis

**Author notes:** **Corresponding author:** Dr. Edmund Hollis II Circuit Repair Laboratory Burke Neurological Institute 785 Mamaroneck Avenue White Plains, New York-10605. **Author contributions:** EH, HM, and YL designed the experiments; HM performed surgery and data analysis. YL, HM and PDG performed mouse behavior. HM, AB and PDG performed data analysis. SA created the illustrations. EH and HM wrote the paper.

## Abstract

Hand and arm manual dexterity is a hallmark of humans and non-human primates. While rodents are less dexterous than primates, they provide powerful models for testing neural circuit function in behavioral output, including dexterous behaviors. In rodents, the single pellet reach task has been used extensively to study both dexterous forelimb motor learning as well as recovery from injury; however, mice exhibit high variability in task acquisition in comparison to rats and a significant percentage fail to learn the task. We have created a recessed version of the task that requires greater dexterity. This subtle modification increases both task difficulty as well as the proportion of mice that show an improvement with training. Furthermore, motor cortex inactivation shows a greater effect on the execution of the recessed forelimb reach task, with distinct effects on reach targeting vs grasping components depending on the timing of inhibitory activation. Kinematic analysis revealed differences in reach targeting upon transient cortical inhibition prior to reach onset. In summary, the recessed single pellet reach task provides a robust assessment of forelimb dexterity in mice and a tool for studying skilled motor acquisition and execution.

## Introduction

Skilled forelimb movements are a defining characteristic of primate motor control and are well developed in rodents as well (Alstermark and Isa 2012, Karl and Whishaw 2013, Whishaw and Karl 2014). Rodents serve as easily accessible and genetically amenable models for studying skilled motor learning and disruption due to injury. The single pellet reach task is frequently used in rat models to study motor learning (Chen, Gilmore et al. 2014, Zemmar, Kast et al. 2015, Bova, Kernodle et al. 2019) or to assess recovery from central nervous system injuries such as spinal cord injury and stroke (Alaverdashvili and Whishaw 2013, Bell, Wolke et al. 2015, Hollis II, Ishiko et al. 2016). Freely moving, head-fixed, and automated versions of this task have been implemented successfully to study motor circuit function (Fenrich, May et al. 2015, Guo, Graves et al. 2015, Wong, Ramanathan et al. 2015, Ellens, Gaidica et al. 2016, Whishaw, Faraji et al. 2018). The task has been characterized in rats with a detailed analysis of reaching and grasping component movements (Whishaw and Pellis 1990, Whishaw, Pellis et al. 1991, Whishaw, Pellis et al. 1992). Deficits in skilled forelimb function can be assessed using this task after pyramidotomy (Whishaw, Piecharka et al. 2003, Piecharka, Kleim et al. 2005), lesions of the corticospinal tract (Carmel, Kim et al. 2010, Hollis II, Ishiko et al. 2016) or red nucleus and rubrospinal tract lesions (Morris, Tosolini et al. 2011, Morris, Vallester et al. 2015).

The execution of skilled, dexterous movements is mediated by motor cortex and its descending projections to the spinal cord (Wang, Liu et al. 2017, Ueno, Nakamura et al. 2018). Topographically organized motor cortex consists of local and corticofugal excitatory projection neurons as well as inhibitory interneurons that delineate region boundaries and modulate the firing of pyramidal neuron ensembles (Tanaka, Tanaka et al. 2011, Kaneko 2013). Disruption of the excitatory/inhibitory balance by injection of GABA agonists shifts the topographic boundaries of motor representations that are shaped by development and refined during motor learning (Jacobs and Donoghue 1991, Galea and Darian-Smith 1995, Kleim, Barbay et al. 1998, Young, Vuong et al. 2012). Motor learning depends on the plasticity of motor networks as attenuation of cortical plasticity mechanisms impairs skilled task acquisition (Li and Hollis 2017).

A major limitation in using the skilled reach task to assess skilled motor learning in mice is that many animals fail to successfully learn the task, exhibiting an essentially flat learning curve (Chen, Gilmore et al. 2014). In this paper, we describe a recessed version of the skilled reach task in which mice have to retrieve a food pellet from a concave depression, similar to the Kluver board test. The Kluver board behavioral test uses wells of different sizes to measure manual dexterity in non-human primates. Monkeys use the thumb and index finger to form a precision grip in order to retrieve a raisin from the wells (Murata, Higo et al. 2008, Qi, Gharbawie et al. 2013, Sugiyama, Higo et al. 2013). The prehension ability of mice to reach into a recession and grasp food pellets is a hallmark of corticospinal function (Gu, Kalambogias et al. 2017). The recessed forelimb reach task proved to be of greater difficulty for mice. Furthermore, we more reliably demonstrated a role of the motor cortex in executing this learned dexterous behavior and found that optogenetic silencing of motor cortex consistently impaired success on the recessed skilled reach task. Kinematic analysis of forelimb reach using DeepLabCut showed altered modulation of reach parameters during cortical silencing. The recessed skilled reach task is a simple, but powerful modified version of a standard behavioral task that will provide a critical tool for applying the genetic tools of mice to motor learning and injury studies heretofore performed in rats.

## Materials and Methods

All procedures were approved by the Burke Neurological Institute/Weill Cornell Medicine Institutional Animal Care and Use Committee. Animals were housed on 12-hour light and dark cycle and behavioral training was performed at consistent time periods.

### Animals

Behavioral experiments were performed on 22 adult mice on the C57BL6J background. Optogenetic silencing and behavior experiments were done in 10 adult C57BL6J mice and 10 adult *Pvalb-Cre::Ai14 Rosa-LSL-tdTomato* mice (The Jackson Laboratory).

### Optogenetic silencing of the motor cortex

Adult C57BL6J or *Pvalb-Cre::Ai14* mice were deeply anesthetized with isoflurane using a SomnoSuite anesthetic system (Kent Scientific), until unresponsive to toe and tail pinch. The skin over the skull was shaved and cleaned with three alternating washes with povidone-iodine solution and 70% ethanol before incision. Small craniotomies were made over forelimb or hindlimb motor cortex for injections through intact dura. A 1:1 mixture of Cre dependent AAV2-Ef1a-DIO-IC++-EYFP (Berndt et al., 2016; 5.9×10^12^ VP/ml) and AAV1 Cre (2.94×10^13^ VP/ml) was injected into C57BL6/J mice. *Pvalb-Cre::Ai14* mice were transduced with Cre-dependent AAV encoding channelrhodopsin (AAV2-Ef1a-DIO-hChR2(H134R)-EYFP (Add gene; 4.2×10^12^ VP/ml). Viruses were injected into either 4 sites in forelimb (AP/ML: 0/1.8, 0/2.4, 1.0/1.5, 1/2.0 mm) or hindlimb (AP/ML: −1.2/1, −1.2/1.5, −1.5/1.5 mm) motor cortex. 300 nl of virus solution was injected at a depth of 500 μm at a rate of 40 nl/min. Following injection, the pipette was left in place for five minutes before withdrawal from the brain. A 2.5 mm stainless steel cannula with guide (Thorlabs) was then placed above the cortical surface using a cannula holder and glued in place using cyanoacrylate glue and C&B-Metabond dental adhesive (Parkell). Mice were placed on a heating pad until alert, then moved to home cages.

### Skilled reach behavior

The single pellet box used for testing and training the animals was made from 3 mm thick transparent cast acrylic. The box measured 8.4 cm width × 19.7 cm depth × 19.7 cm height. A sliding door (15.1 cm width × 19.2 cm height × 3 mm thick with two 1 cm wide slits set at 1.6 cm from each side) was positioned with an opening on the right or the left side based on the dominant paw of the mouse. Food pellets were placed on a white block (8 cm length × 6 cm width × 0.9 cm height) with one small indentation (< 0.5 mm depth) for standard forelimb reach, or with a small recession (2 mm depth, 6 mm diameter) for recessed forelimb reach.

Two days before the beginning of the training, each mouse was weighed and put on food restriction. Mice were weighed daily and, after the training, a measured and specific amount of food was given to each mouse to maintain body weight at or above 80% of pre-restriction weight. Animals were trained daily on the standard or recessed versions of the forelimb reach task for a period of two weeks. Animals extended a forelimb through the vertical slot in the front sliding door and over a small gap to retrieve food pellets. 20 mg food pellets (Dustless Precision Pellets Rodent, Purified Chocolate Flavor, Bio-Serv, Flemington, NJ) were placed 1 cm away from the inside of the front wall of the box in the indentation or recession positioned at the medial edge of the slit through which the mouse reached. Mice performed 25 reaches per session, in which the paw extended and made contact with the pellet. Successful reaches were scored when the mouse retrieved and ate the pellet. Success rate was calculated as the number of pellets successfully retrieved divided by the total number of reaches.

Mice for optogenetic experiments were trained with the fiber optic cable attached during the entire duration of the behavior. Optogenetic silencing was performed with blue light (473 nm LED, 7.5 mW/m^2^) for two days with LED ON or LED OFF with 25 reaches per session. The order of LED ON and OFF sessions was randomized each day. ChR2 activation of parvalbumin interneurons in Pvalb-Cre mice was performed after two weeks of training on recessed reach behavior. Parvalbumin neurons were activated using blue light (473 nm LED, 7.5 mW/m^2^) for 25 trials of LED ON and LED OFF sessions for four days. The mice were perfused after finishing the silencing or activation experiments.

### Behavioral video analysis

Videos were recorded using a Basler Ace camera (acA1440-220um) at 33 frames per second. Kinematics of individual reaches were analyzed using frame by frame video analysis of recorded behavior and markerless pose estimation. The total number of reach attempts was counted from video recordings. Videos were imported into the video analysis and modeling software tool Tracker (version 5.1.4). Rotational angles of pronation, supination 1, and supination 2 were measured (Carmel et al., 2010) using the protractor function for a minimum of five reaches per session and compared between groups for LED ON and OFF sessions. The machine learning algorithm, DeepLabCut was used to track the pellet and digits 2-5 during reach to reconstruct the trajectory of the forelimb during extension and pellet retrieval in both LED ON and OFF sessions (Mathis, Mamidanna et al. 2018).

In DeepLabCut, four videos were used to compose the training set. From each video, 40 frames were extracted where digits 2-5 and the pellet were labelled. These frames were used to refine the pre-trained network (ResNet) to predict features of interest in unseen videos and generate x- and y-coordinates for each label throughout the video. These coordinates were imported into Matlab R2018b along with labeled start and end frames of each reach attempt. A spatial calibration factor was used to convert pixel values to millimeters. Trajectory reconstruction was done by finding the center of mass of the paw based on digit coordinates and calculating the linear best fit line of all reach attempts per animal per session. Left paw trajectories were reflected using the stationary pellet as the line of reflection. Trajectories of mean reach attempts were compared between LED ON and LED OFF conditions by calculating the mean absolute distance between the two best-fit lines.

### Perfusions and histology

Mice were deeply anesthetized with ketamine-xylazine cocktail, transcardially perfused with ice-cold PBS followed by 4% paraformaldehyde in PBS, and brains were post-fixed overnight in 4% paraformaldehyde. Cortical hemispheres were isolated from underlying structures and were flattened between glass slides and cryoprotected with 30% sucrose. The flattened cortex was sectioned on a sliding microtome (Leica) at 50 um thickness. Alternate series of sections were mounted onto glass slides, dried and coverslipped with Cytoseal 60 and stored at 4°C. The other series of sections were reacted for cytochrome oxidase (CO) in a water bath at 42°C for 1.5-2 hours, checked for staining intensity and mounted onto glass slides (Wong-Riley 1979, Jain, Diener et al. 2003). Dried sections were coverslipped with Cytoseal 60 and stored at room temperature.

### Fluorescence microscopy and reconstruction of injection sites

The cortices of *Pvalb-Cre::Ai14* mice (n=5) and C57BL6J mice (n=10) were analyzed for the spread of virus injections in the motor and somatosensory areas. The spread of gene expression was outlined using a fluorescent microscope installed with Neurolucida software (version 9, MBF Bioscience). EMF files were exported from Neurolucida into Canvas x Draw software (version 6, Canvas GFX, Inc) and sections were aligned using blood vessels and tissue landmarks. Outlines of injection sites from individual mice were overlaid on a section of the flattened hemisphere stained with cytochrome oxidase (Mohammed and Jain 2016).

### Statistical analysis

Behavior data were analyzed in Prism version 8 (GraphPad Software, LLC). Paired tests were performed on parametric data using Prism, as indicated in the results. Individual groups were tested and analyzed by an investigator blinded to grouping.

## Results

### Mice more reliably improve with training on a recessed version of the forelimb reach task

Seven mice were trained on a standard forelimb reach task in which the food pellet is placed in a small indentation that provides for consistent placement. On this standard task, the initial success rate was 27 ± 5% (mean ± s.e.m.), while after two weeks of training this had only increased to 34 ± 4% (Fig. 1a,b). The resulting increase was not statistically significant (P = 0.153; paired, two-tailed t-test). Fifteen separate mice were trained on a modified version of the forelimb reach task in which the pellet is placed in a small recession measuring 2 mm deep and 6 mm in diameter, situated 1 cm from the interior wall of the box. Mice had an initial success rate of 17 ± 3% on the recessed forelimb reach task. After two weeks of training, they increased to 41 ± 3% on average, a highly significant improvement (P < 0.0001; paired, two-tailed t-test). The recessed reach group exhibited a steeper learning curve, with a lower initial performance. Previously it has been described that there are mice that fail to learn the task, either starting at a high-proficiency (“over-shaped”) or maintaining a poor performance throughout training (“non-learners”) (Chen, Gilmore et al. 2014). These mice do not improve by more than 15% of their initial performance. We found that 3 of 7 mice (43%) trained on the standard forelimb reach task failed to improve by more than 15% over the course of two weeks (Fig. 1). These non-learners succeeded in retrieving 37 ± 4% of the food pellets on the first day of training, and an average of only 28 ± 5% over the last three days. Four mice did manage to learn the task and improve their performance by more than 15%, starting at 19 ± 7% on day 1 and ending at 41 ± 6% averaged over the last 3 days. In contrast, 93% of the mice trained on the recessed forelimb reach task demonstrated improvements with training, increasing from 16 ± 3% success on day 1, to 43 ± 3% averaged over days 12-14. Only one mouse tested on the recessed version of the task failed to significantly improve, starting at 32% and ending at a 35% success rate.

**Figure 1.**
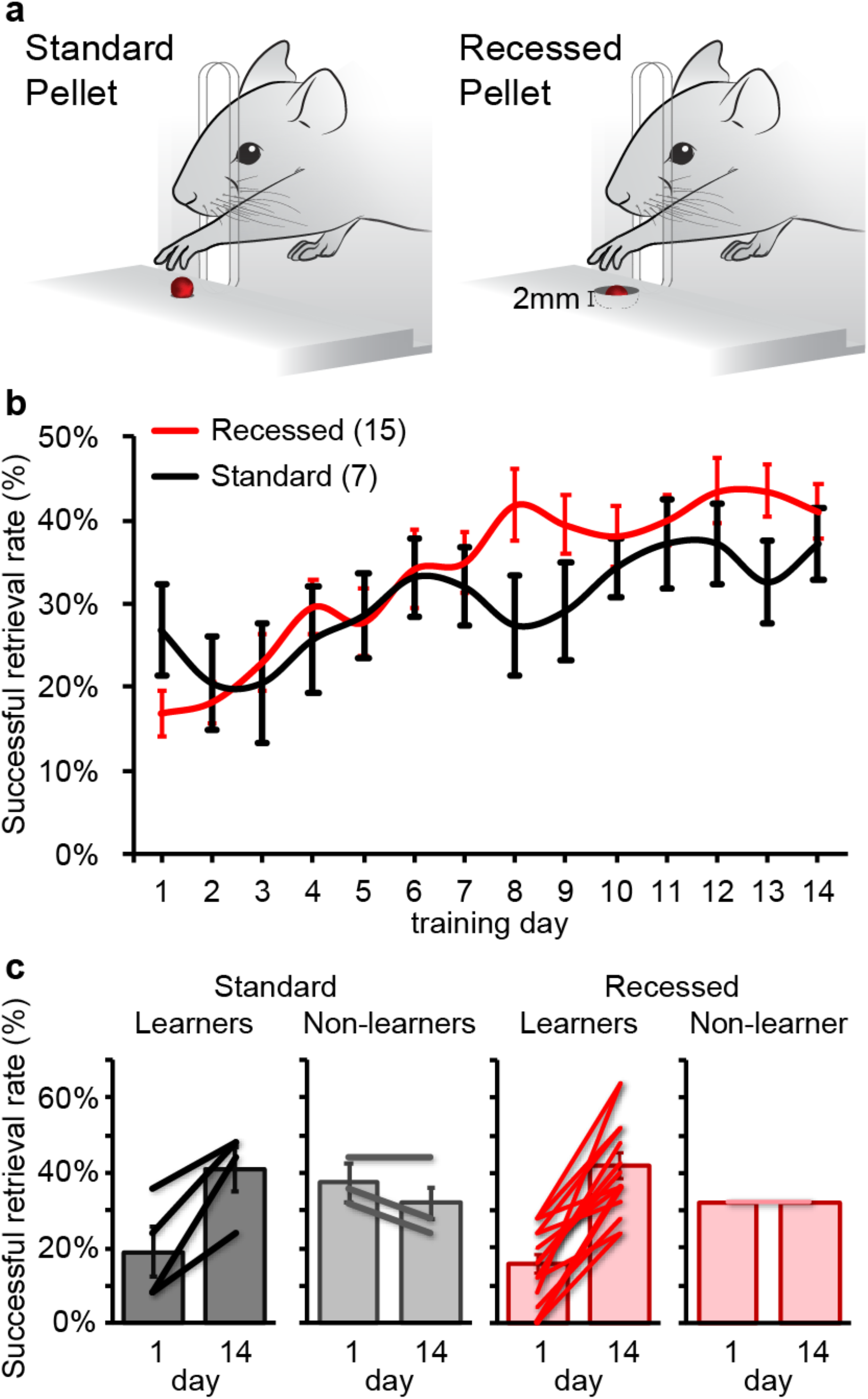
Mice exhibit more consistent learning on the recessed forelimb reach task. (a) Illustration of the standard and recessed forelimb reach tasks. (b) Successful retrieval rate significantly improves over 2 weeks of behavioral training on the recessed version compared to the standard task (repeated measures ANOVA *P* = 0.04). (c) The majority of mice exhibit learning over the course of training on the recessed task (14/15), in contrast to the standard task (4/7). Data presented as mean ± sem.

### Motor cortex silencing elicits greater effects on recessed forelimb reach performance

After establishing forepaw preference, forelimb or hindlimb motor cortex in adult C57BL6J mice was transduced to express the inhibitory chloride conducting channelrhodopsin variant iC++ (Berndt, Lee et al. 2016). Mice were then trained on the standard version of the skilled reach behavior for two weeks with a fiber optic cable attached to a head-mounted cannula. Following training, mice were tested on two consecutive days with two sessions consisting of 25 reaches, one with 473 nm LED ON and one with the LED OFF. The sequence of LED ON and OFF sessions were alternated for each mouse. LED activation was triggered during the paw advance phase by an IR sensor located outside of the reach chamber (Fig. 2). The average success rate during the activation of iC++ in forelimb motor cortex dropped from 63.6 ± 2.9% during LED OFF sessions to 51.2 ± 4.5% during LED ON sessions, a non-significant difference of 12.4% (paired, two-tailed t-test *P* = 0.16). Activation of iC++ expressed in hindlimb motor cortex resulted in no measurable impairment on the standard forelimb reach, with an average success rate of 58 ± 4% during LED OFF sessions and 55.2 ± 5.7% during LED ON sessions, a difference of less than 3% (paired, two-tailed t-test *P* = 0.21).

**Figure 2.**
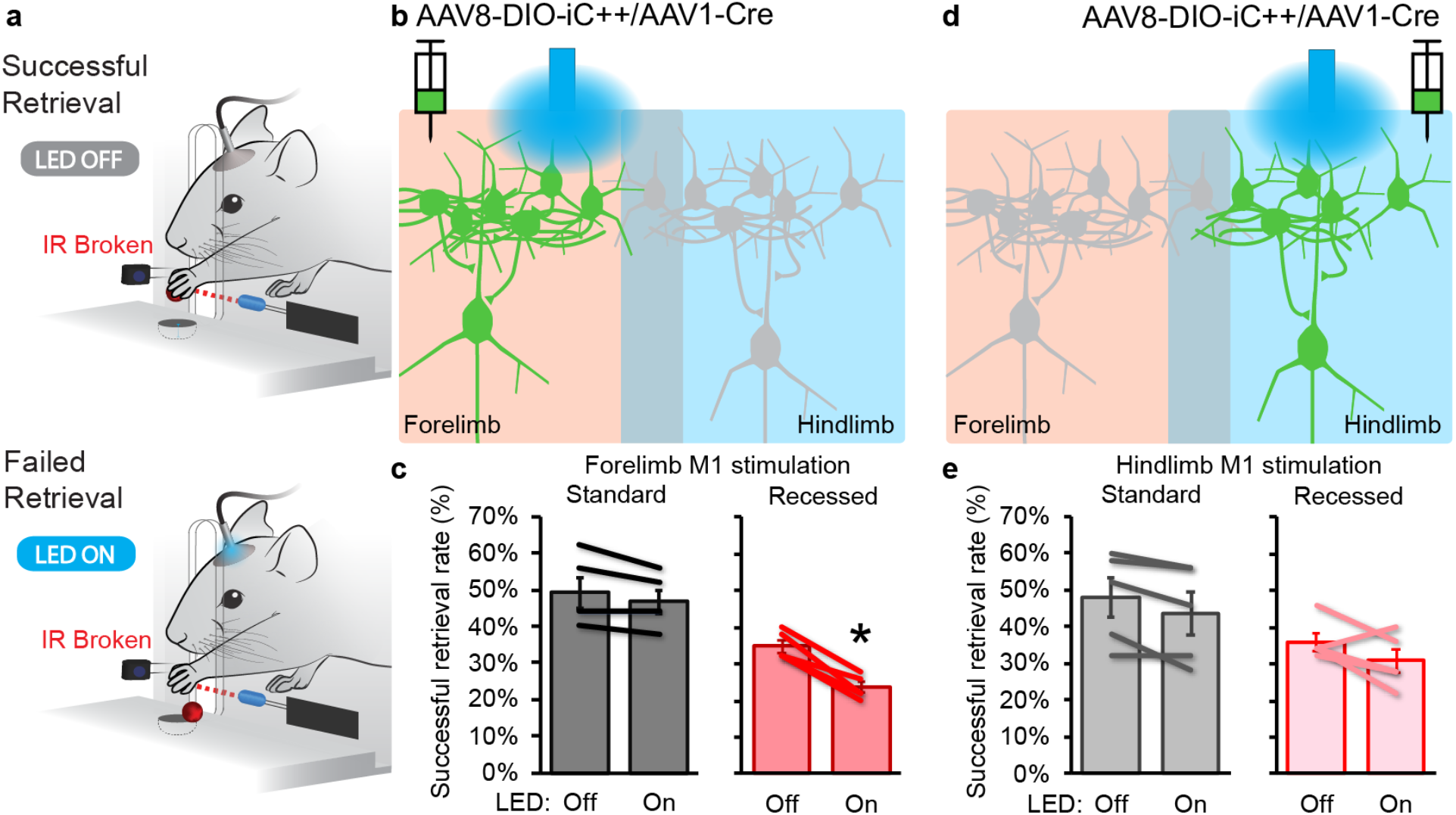
Silencing forelimb motor cortex impairs dexterous motor performance on the recessed forelimb reach task. (a) Illustration of IR LED-sensor triggered optogenetic silencing. (b) Illustration of local cortical transduction for forelimb motor cortex silencing through chloride-conducting iC++ activation. (c) 473 nm LED stimulation of iC++ transduced forelimb motor cortex disrupts recessed, but not standard forelimb reach (paired, two-tailed t-test *P* = 0.002). (d,e) Silencing of iC++ transduced hindlimb cortex did not significantly affect performance on either task. Data presented as mean ± sem; *n* = 5/group.

In contrast to the effects of cortical silencing during the standard forelimb reach, silencing forelimb motor cortex significantly impaired the execution of the dexterous recessed forelimb reach. After iC++ testing sessions on the standard reach, mice were re-trained on the recessed reach over a period of one week. Following training, mice were tested on two consecutive days as above. Silencing of forelimb motor cortex by iC++ activation resulted in an 11% drop in success rate, from 34.8 ± 1.7% during LED OFF sessions to 23.6 ± 1.5% during LED ON sessions (paired, two-tailed t-test *P* = 0.002). Silencing of hindlimb areas had little effect on recessed reach performance, with average success rate dropping by 5% from 36.0 ± 2.5% during LED OFF sessions to 30.8 ± 3.2% during LED ON sessions (paired, two-tailed t-test *P* = 0.17). The ratio of LED ON/LED OFF success rate on the recessed forelimb reach was 68.1 ± 4.0% for forelimb area silencing and 86.1 ± 8.7% for hindlimb area silencing (Fig. 2).

### Motor cortex silencing alters reach targeting but does not alter the biomechanics of successful forelimb reach grasping

In addition to reach success rate, behavioral videos were analyzed for variability in the total number of attempts needed to complete 25 reaches. Reaches consist of attempts during which the pellet is touched by the forepaw; however, cortical injury can disrupt aiming during the reaching movement. In order to determine whether transient cortical silencing affects the trajectory of forelimb reach behavior, we quantified the number of attempted reaches during which the forepaw failed to make contact with the food pellet. The total number of attempts needed to make 25 reaches during silencing of the forelimb motor cortex exhibited a moderate increase, with a 20.0 ± 7.0% increase in total reach attempts during the standard forelimb reach (ratio paired, two-tailed t-test *P* = 0.03) as well as during the recessed forelimb reach (18.9 ± 8.3%, ratio paired, two-tailed t-test *P* = 0.04). In contrast, silencing of hindlimb motor cortex did not significantly alter the number of off-target reach attempts during either the standard (12.3 ± 12.2%, ratio paired, two-tailed t-test *P* = 0.52) or recessed (11.3 ± 6.3%, ratio paired, two-tailed t-test *P* = 0.17) forelimb reach. (Supp. Fig. 1).

The stereotypic forelimb reach movement can be broken down into individual components. In addition to impairment of overall success, damage to the motor cortex induces deficits in execution of some of these components, primarily those related to paw rotation (Whishaw and Pellis 1990, Whishaw, Pellis et al. 1991). The rotational components of individual pellet grasps (pronation, supination 1, and supination 2) from a minimum of 5 successful reaches were analyzed frame by frame from video recordings of both standard and recessed forelimb reach. Unlike the effects previously observed after cortical injury, transient silencing of forelimb or hindlimb motor cortex did not significantly alter forepaw pronation or supination components of forelimb reach behavior. On the standard forelimb reach task, the average difference in angles between LED ON and LED OFF trials (ON-OFF) when silencing the forelimb motor cortex was - 0.8 ± 1.8 degrees during pronation, 1.8 ± 1.8 degrees during supination 1, and 5.6 ± 3.8 degrees during supination 2. Likewise, silencing of forelimb motor cortex during recessed reach did not significantly affect rotational movements of the forepaw. The average ON-OFF difference in angles during recessed reach were −2.5 ± 2.8 degrees during pronation, 4.1 ± 4.1 degrees during supination 1, and 4.3 ± 4.1 degrees during supination 2. Silencing of hindlimb motor areas had no significant effect on the execution of rotational components of either standard or recessed forelimb reach. The average ON-OFF angle differences during standard forelimb reach were −0.4 ± 2.2 degrees during pronation, 5.5 ± 2.5 degrees during supination 1, and 1.2 ± 2.2 degrees during supination 2, while the values for ON-OFF angle differences during recessed reach were 2.7 ± 0.6 degrees during pronation, −2.9 ± 3.2 degrees during supination 1, and −2.5 ± 3.7 degrees during supination 2.

### Activation of parvalbumin-expressing GABAergic neurons disrupts execution of trained dexterous pellet reach

Following our studies using indiscriminate silencing of both excitatory and inhibitory cortical neurons, we specifically inhibited the activity of glutamatergic cortical neurons by stimulating channelrhodopsin expressed in parvalbumin-expressing GABAergic neurons within forelimb or hindlimb regions of motor cortex. Parvalbumin neurons are the largest population of interneurons in the neocortex (Tremblay, Lee et al. 2016). They regulate pyramidal neuron ensemble timing through extensively branched axons and robust inhibitory inputs to pyramidal cell somata and proximal dendrites or axon initial segments (Hu, Gan et al. 2014). Hyperpolarizing GABA release close to the site of action potential generation leads to robust inhibition of pyramidal cell output. *Pvalb-Cre::Ai14* mice transduced with Cre-dependent AAV encoding channelrhodopsin in either forelimb or hindlimb motor cortex were trained on the recessed forelimb reach task for two weeks. Following training, channelrhodopsin-expressing parvalbumin neurons were activated using two distinct paradigms during task execution. First, the LED was manually turned ON during the preparatory phase of the reach, in order to mimic spontaneous forelimb motor cortex parvalbumin neuron activation that occurs prior to reach onset (Estebanez, Hoffmann et al. 2017). The LED was turned on as the mouse approached the front side of the box and kept ON throughout the reach. The second testing paradigm used an Arduino based closed loop system activated by an IR sensor. As mice advanced their paws towards the food pellet, interruption of an IR beam path triggered the LED ON until the paw was retracted.

Activation of parvalbumin neurons in the motor cortex during the preparatory phase of the reach elicited no major effect on the success of the reach. In forelimb motor cortex transduced mice, the average success rate was 34.4 ± 4.4% during the LED OFF sessions and 36.3 ± 6.3% during sessions with activation of forelimb parvalbumin neurons prior to reach onset, an average difference in success rates (ON-OFF) of 1.9+-2.5% (paired, two-tailed t-test *P* = 0.62; Fig. 3c). Similarly, silencing hindlimb cortex during the preparatory phase had no significant effect on reach performance. The average success rate during LED OFF sessions was 35.7 ± 0.33%, and 41.0 ± 2.1% when LED ON sessions began during reach onset. The average difference in success rate with hindlimb preparatory parvalbumin activation (ON-OFF) was 5.3 ± 2.2% (paired, two-tailed t-test *P* = 0.09; Fig. 3c).

**Figure 3.**
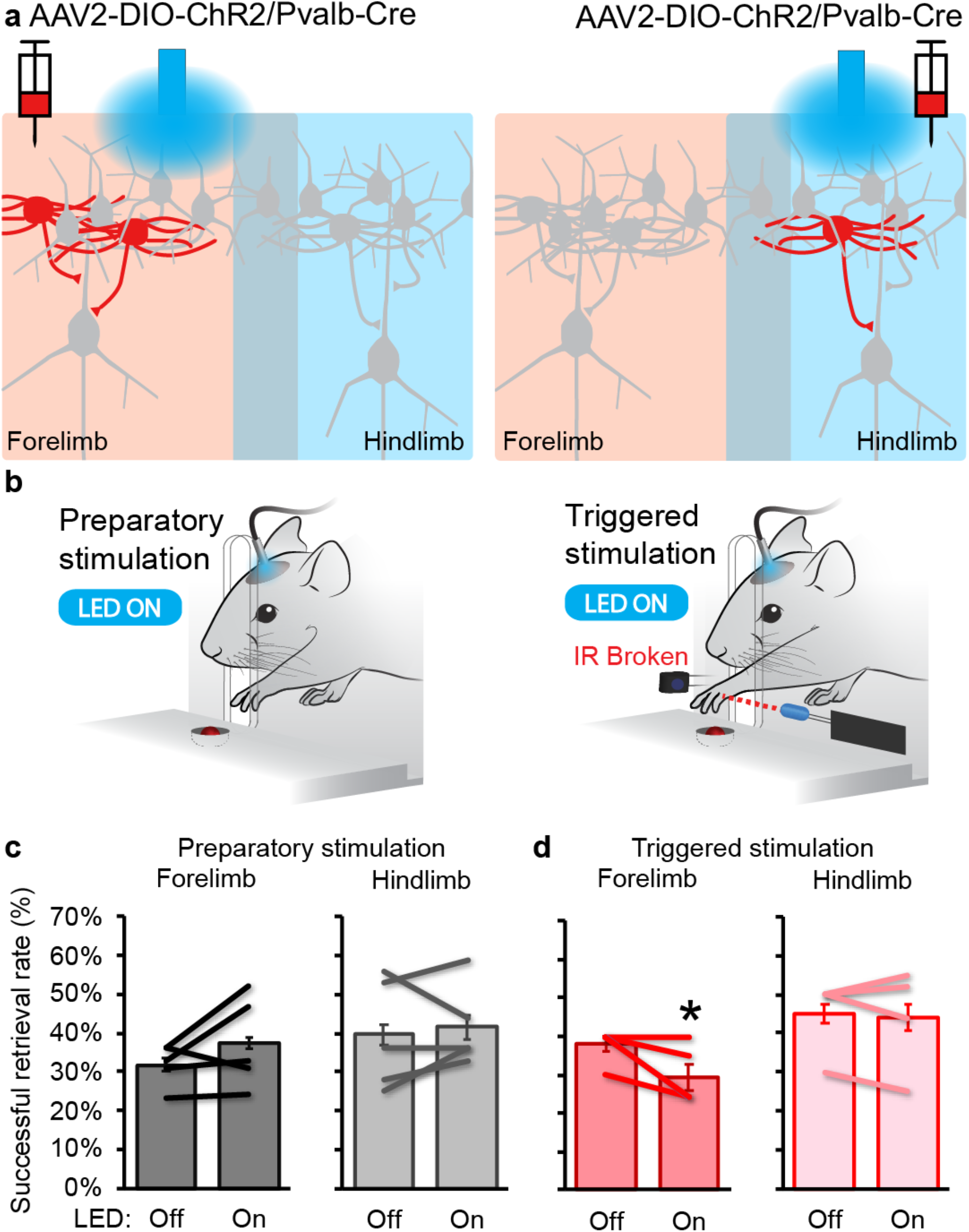
Timing of parvalbumin neuron activation in forelimb motor cortex determines reach impairment. (a) Illustration of local parvalbumin neuron transduction in forelimb and hindlimb motor cortex with AAV2-DIO-ChR2 (indicated in red). (b) Illustration of 473 nm LED stimulation of Pvalb-ChR2 before reach initiation (left) or in response to IR LED-sensor trigger during reach. (c) Activation of Pvalb neurons prior to reach onset did not alter successful retrieval upon contact with the pellet. (d) Activation of Pvalb neurons in forelimb cortex immediately prior to pellet touch impairs successful retrieval (paired, two-tailed t-test *P* = 0.05). Data presented as mean ± sem; *n* = 5/group (except *n* = 4 in triggered hindlimb group).

In contrast to the effects of stimulating parvalbumin neurons during the preparatory phase, increasing parvalbumin activity in forelimb motor cortex during execution of the reach impaired performance of the recessed forelimb reach task. The average success rate during LED OFF trials was 37.7 ± 3% in mice expressing channelrhodopsin in forelimb motor cortex parvalbumin neurons. During sessions when the IR sensor triggered LED activation of parvalbumin neurons during reach, the success rate dropped to 30.7 ± 4.1%, an average difference in success rates of −7.0 ± 1.2% (paired, two-tailed t-test *P* = 0.05, Fig. 3d). Activation of parvalbumin neurons in hindlimb regions of motor cortex during the reach elicited no significant impairment in this forelimb behavior. Success of hindlimb transduced mice was 45.0 ± 3.2% during LED OFF sessions and 44.0 ± 2.5% during LED ON sessions (paired, two-tailed t-test *P* = 0.73; Fig. 3d)

### Activation of parvalbumin neurons prior to reach onset alters reach targeting, but does not alter the biomechanics of successful forelimb reach grasping

Similar to iC++ silencing experiments, we also analyzed whether parvalbumin activation affected the targeting phase of reaching by quantifying the number of attempts necessary to perform 25 reaches. Only activation of forelimb motor cortex parvalbumin neurons during the preparatory phase disrupted targeting. Total reach attempts increased by 18.6 ± 4.6% when these neurons were activated prior to reach initiation (ratio paired, two-tailed t-test *P* = 0.02). Activation of the same forelimb motor cortex parvalbumin neurons during the execution of the movement did not affect targeting, with highly variable average increase in total attempts of 5.6 ± 11.1%. Likewise, activating hindlimb motor cortex parvalbumin neurons either during the preparatory phase or the execution phase did not increase the number of total reach attempts (1.3 ± 3.3% during preparatory, 9.1 ± 5.2 during execution).

Similar to iC++ silencing of motor cortex, activation of neither forelimb nor hindlimb motor cortex parvalbumin neurons significantly disrupted the rotational components of learned reaching behavior. This was true for stimulation during both the preparatory and execution phases of the reach. The average ON-OFF difference in angles when activating forelimb motor cortex parvalbumin neurons during the preparatory phase was 3.7 ± 5.1 degrees during pronation, −6.0 ± 3.8 degrees during supination 1, and 0.4 ± 1.3 degrees during supination 2. When forelimb motor cortex parvalbumin activation occurred during reach execution, the ON-OFF difference in angles was 5.0 ± 1.8 degrees during pronation, 11.2 ± 9.7 degrees during supination 1, and 4.6 ± 8.4 degrees during supination 2.

Similarly, there was no significant difference on rotational angles during activation of hindlimb motor cortex parvalbumin neurons. The average ON-OFF angle differences when stimulated during the preparatory phase were −0.4 ± 2.2 degrees during pronation, 5.5 ± 2.5 degrees during supination 1, and 1.2 ± 2.2 degrees during supination 2; during the execution phase, the average ON-OFF angle differences were 1.9 ± 2.5 degrees during pronation, 2.9 ± 4.0 degrees during supination 1, and −2.8 ± 3.4 degrees during supination 2.

### Silencing prior to reach onset exerts larger effects on reach targeting than silencing during execution

DeepLabCut markerless kinematic analysis was conducted on reaches performed by mice expressing channelrhodopsin in parvalbumin-expressing neurons. The difference between mean, best-fit linear trajectories during LED on and off sessions was found to be significantly higher in mice with forelimb motor cortex silencing during the preparatory phase than when silencing was triggered during reach execution (*n* = 4; paired, two-tailed t-test *P* = 0.03; Fig 4d). Silencing of hindlimb cortex during the preparatory phase showed a less consistent disruption of reach trajectory compared to silencing during reach execution (*n* = 4; paired, two-tailed t-test *P* = 0.14; Fig 4g).

**Figure 4.**
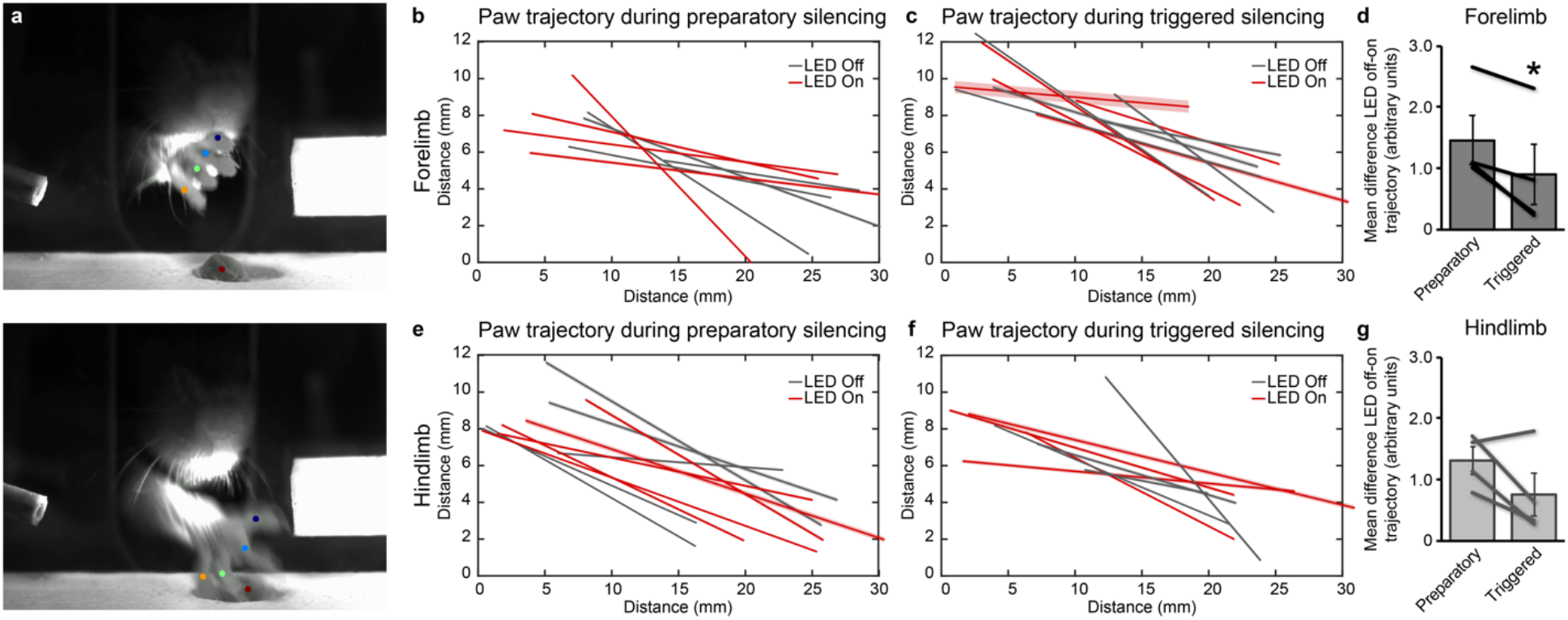
Motor cortex silencing prior to reach onset disrupts reach trajectory. (a) Example of forepaw labeling used for the machine learning algorithm DeepLabCut kinematic analysis. (b,c) Forepaw trajectories during recessed reach when activating forelimb region ChR2-expressing parvalbumin neurons (b) prior to, or (c) during forelimb reach. (d) Silencing of forelimb cortex prior to reach onset resulted in a greater disruption of forepaw trajectory than silencing during reach (paired, two-tailed t-test *P* = 0.03). (e,f) Forepaw trajectories when hindlimb ChR2-expressing parvalbumin neurons were activated (e) prior to, or (f) during forelimb reach. (g) Disruption of forepaw trajectory during silencing of hindlimb cortex. Data presented as mean ± sem; *n* = 4/group.

## Discussion

In this paper we describe a modified version of the forelimb reach task for mice as well as the effects of temporally regulated motor cortex silencing on behavior. We demonstrated that our recessed forelimb reach task provides a more powerful tool to study dexterous, forelimb, motor control in mice than previous iterations. During training, a lower initial performance on the recessed task than on the standard reach indicated that the recessed pellet was more difficult to successfully retrieve. This initial performance provided a baseline that allowed for a clear learning curve over two weeks of training. In contrast, a large proportion of mice trained on the standard forelimb reach task showed a limited improvement with training. We also found that transient silencing of motor cortex more reliably disrupted execution of the trained performance on the recessed version of the task, implicating a key role for motor cortex in modulating trained dexterous movements. The timing of cortical silencing was of critical importance, with silencing prior to reach initiation affecting targeting, while silencing during the reach disrupted grasp and pellet retrieval. Notably, though interrupting cortical activity affected overall success rates, it did not measurably alter the rotational kinetic components of the successful reaches. Our data indicates that motor cortex activity is not required for the engrained, ballistic components of the trained movement, but is rather likely a critical mechanism for adjusting the choreographed execution of complex, patterned, grasping movements.

### Role of motor cortex in the execution of skilled forelimb movements

Rodents have been used to test the contribution of motor cortex and corticospinal projections in the execution of skilled forelimb reach behavior in studies employing cortical ablation, ischemic motor cortex stroke, or neonatal hemi-decortications (Castro 1972, Barth, Jones et al. 1990, Gharbawie, Gonzalez et al. 2005, Takahashi, Vattanajun et al. 2009, Umeda and Isa 2011, Kawai, Markman et al. 2015). The plasticity of motor cortex that supports motor learning is also required for restoring function after brain or spinal cord injury in mammals (Kambi, Tandon et al. 2011, Mohammed and Hollis 2018). Only recently have powerful transgenic mouse models been used in the study of learned motor control (Azim, Jiang et al. 2014, Hollis II, Ishiko et al. 2016, Wang, Liu et al. 2017, Ueno, Nakamura et al. 2018, Sathyamurthy, Barik et al. 2020).

In direct comparisons between the standard and recessed version of the forelimb reach task, we found that optogenetic silencing of the cortex disrupted forelimb reach targeting and increased the number of failed reach attempts. When mice did successfully reach the pellet, cortical silencing only affected success on the recessed version of the task, indicating that activity in forelimb motor areas is required for execution of the dexterous aspects of forepaw grasping. Regional silencing or photoinhibition of caudal forelimb area (CFA) in a directional joystick task impairs motor execution and fine motor control (Morandell and Huber 2017) or peak trajectory speed and outward reaching, attenuating reach trajectory (Bollu 2018, Bollu, Whitehead et al. 2019). Microstimulation and optogenetic mapping have demonstrated strong distal motor (wrist and digit) representations in CFA, with RFA largely evoking wrist and elbow movements (Ramanathan, Conner et al. 2006, Tennant, Adkins et al. 2011, Hollis II, Ishiko et al. 2016). CFA exhibits extensive remapping with skilled behavioral training whereas RFA does not show such changes (Kleim, Barbay et al. 1998). Indeed, silencing RFA shows only minimal impairments on a directional joystick task (Bollu 2018). Our study focused on silencing the neuronal populations in the forelimb or hindlimb motor cortex, specifically CFA and caudal hindlimb area (CHA) (Mohammed and Jain 2014, Mohammed and Jain 2016). These caudal motor areas overlap with the somatosensory representations of the forepaw and hindpaw in somatosensory cortex (S1) (Halley, Baldwin et al. 2020). Hence our approach might have silenced the motor cortex and part of the somatosensory cortex. Parvalbumin neurons in the same areas were activated during the channelrhodopsin experiments. This is evident from histological analysis of AAV injection sites performed on flattened cortical sections (Sup Fig 5).

Others have demonstrated that broadly silencing motor cortex through activation of multiple classes of GABAergic neurons halts the initiation or impairs the execution and trajectory of trained forelimb reach movements (Guo, Graves et al. 2015, Galiñanes, Bonardi et al. 2018). Release of cortical inhibition then lead to a resumption of the trained movement (Guo, Graves et al. 2015), which is reminiscent of the large-scale inhibition observed in preparation for a cued movement (Hasegawa, Majima et al. 2017). In contrast, we selectively activated parvalbumin neurons, the most abundant class of GABAergic cortical neurons. Activation of parvalbumin neurons disrupted the successful execution of our prehension task, as it does in other forelimb motor tasks (Hwang, Dahlen et al. 2019), but did not elicit the robust inhibition of movement initiation or execution observed with non-selective GABAergic activation. Our results suggest that motor cortex parvalbumin neurons may be actively involved in shaping the fine adjustments of prehension, but not necessarily sufficient to interrupt the initiation of movement. Several other subtypes of inhibitory neurons present in the motor cortex (somatostatin, VIP, etc.) likely have significant roles in the modulation of skilled movements.

While it is well-established that motor cortex is a critical center for dexterous motor control, the role of motor cortex in executing learned motor skills diminishes with training. Training on a temporally patterned lever press behavior depends on motor cortex, while execution of the same task after training to proficiency is unaffected by complete ablation of the motor cortex (Kawai, Markman et al. 2015). This is similar to other observations where parvalbumin activation early in training (days 1-9) severely impaired successful execution of a learned lever press task, while motor cortex silencing after extensive training (days 61-69) elicited no significant effect on the behavior (Hwang, Dahlen et al. 2019). Silencing during the middle stages (days 19-26) drove an intermediate impairment (Hwang, Dahlen et al. 2019), at a time on the training schedule similar to what we employed here.

### Timing of parvalbumin activity in skilled forelimb movements

As broad iC++ expression drives chloride influx induced hyperpolarization across a mixed population of excitatory and inhibitory neurons, we also demonstrated how local activation of parvalbumin-expressing GABAergic inhibitory neurons affects recessed forelimb reach. By driving activity of parvalbumin neurons during different phases of the reach, we were able to dissociate the effects on reach targeting from the grasp components required for reach success. Parvalbumin neurons have been found to increase activity prior to the onset of a simple head-fixed reaching task, firing before the pyramidal neurons that initiate movement (Estebanez, Hoffmann et al. 2017). As parvalbumin firing rate was found to gate the amplitude of reach movements (Estebanez, Hoffmann et al. 2017), it is likely that the increase in parvalbumin firing during the reach preparatory phase in our study is driving the aberrant targeting we observed. The preparatory phase of parvalbumin neuron firing has been proposed to modulate neural networks and prevent aberrant or premature activation of corticofugal neurons. This could explain why parvalbumin activation during the preparatory phase disrupted targeting, but activation during the reach was required to disrupt the subsequent grasp aspects of the reach movement. These neurons may underlie command shaping activity of ongoing movements by balanced or recurrent inhibition (Isomura, Harukuni et al. 2009).

### Role of other brain areas in skilled forelimb movements

The somatosensory cortex plays an important role in movement control, incorporating cutaneous or proprioceptive feedback from the skin or joints. In turn, subregions of the motor cortex use distinct sensory modalities to shape motor output (Friel, Barbay et al. 2005). Forelimb S1 drives the adaptation of motor commands and optogenetic inhibition impairs motor adaptation in mice without affecting learned motor patterns (Mathis, Mathis et al. 2017). This is an ongoing modulation as S1 photoinhibition after a partial adaptation impairs the subsequent adaptation while incorporating the ongoing, adapted motor commands. In our experiments, the proximity of sensory and motor regions in the mouse means that we partially silenced S1. As we targeted CFA for silencing, it could be that this predominantly affected cutaneous feedback upon contact with the food pellet, leading to the reduced success observed on the recessed reach.

The behavioral output observed during skilled movements is a net result of computations in discrete sensorimotor regions in the brain, including M1, S1, thalamus, basal ganglia, cerebellum etc. While M1 activity is a critical mediator of fine motor control, coordinated activity across M1 and dorsolateral striatum arises early in skilled training and silencing of the dorsolateral striatum disrupts the gross, but not fine motor movements (Lemke, Ramanathan et al. 2019). The independence of reach and grasp components can be observed behaviorally (Whishaw, Faraji et al. 2017) and the discrete effects of temporal parvalbumin neuron activation that we observed likely alter these individual components independently. Within the striatum, simple and dexterous trained motor movements depend on the dorsolateral striatum, which receives sensorimotor input from M1. Lesion of dorsolateral striatum, but not dorsomedial striatum, impairs learned motor behaviors (Whishaw, Zeeb et al. 2007, Dhawale, Wolff et al. 2019). Dorsolateral striatum appears to be necessary and sufficient to control simple trained movements on a lever press task in the absence of the motor cortex; however, cortical aspiration leads to increased inter-trial variability (Dhawale, Wolff et al. 2019, Wolff, Ko et al. 2019). Combined with the extended training studies showing a disassociation of M1 from execution of trained motor movements (Hwang, Dahlen et al. 2019), it appears likely that the basal ganglia is sufficient to drive execution and perhaps adaptation of learned motor commands during the later stages of motor learning or retention. Whether this holds true for more dexterous tasks, such as our recessed pellet reach task remains unknown.

## Acknowledgments

Authors would like to thank Francesca Marino, Rachel Garn, and Marwa Soliman for technical support. This work was supported by Burke Foundation and NIH Common Fund DP2 NS106663 to EH. HM and YL received post-doctoral fellowships from NYS DoH SCIRB (C33610GG and C32633GG, respectively).

---

**Supplementary Figure 1.**
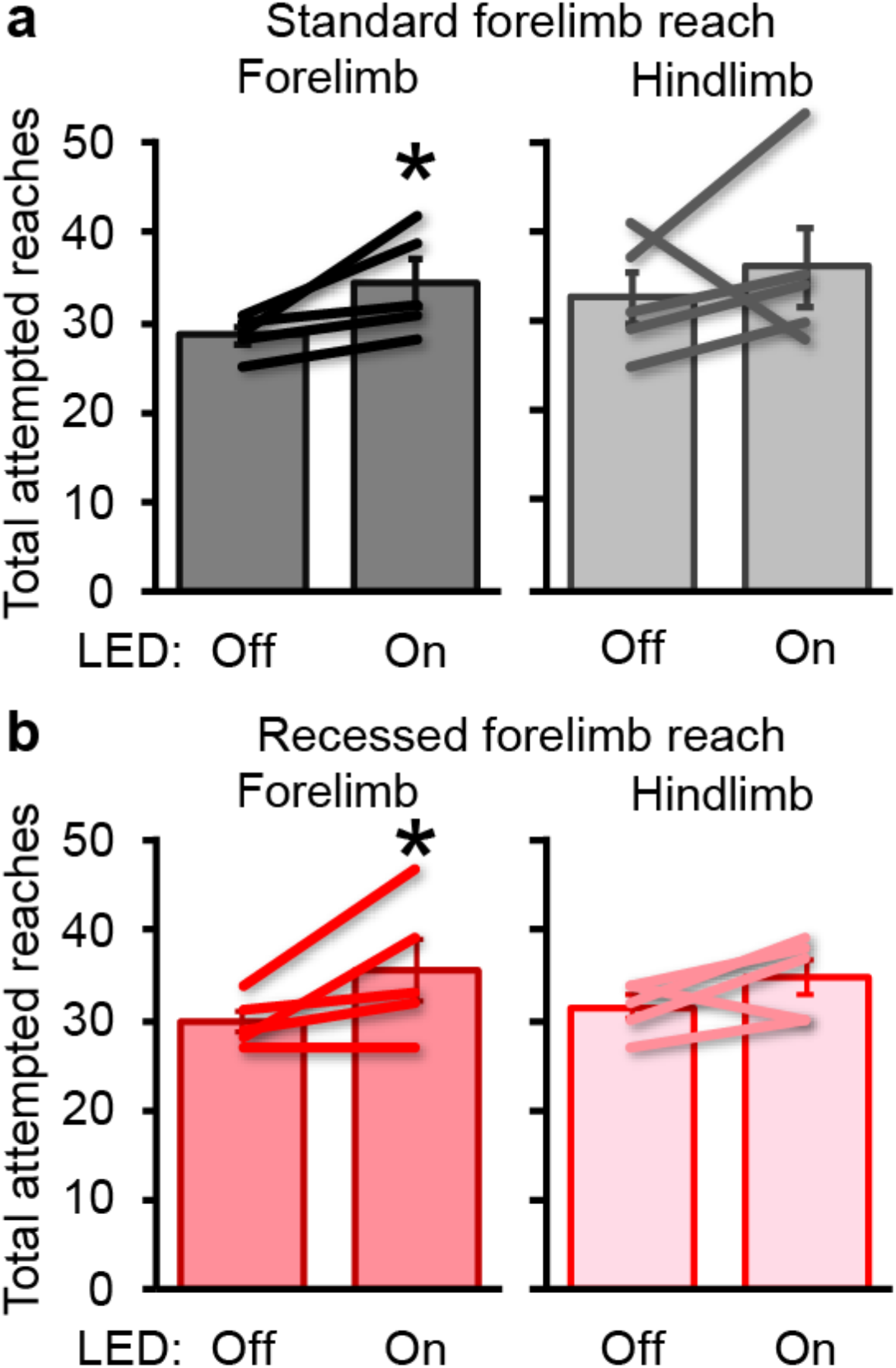
Optogenetic silencing of forelimb motor cortex impairs reach targeting. The total number of attempts needed to complete 25 reaches to the pellet increased when forelimb motor cortex was silenced through iC++ on both standard (a) and recessed (b) forelimb reach tasks (ratio paired, two-tailed t-test *P* < 0.05). Data presented as mean ± sem; *n* = 5/group.

**Supplementary Figure 2.**
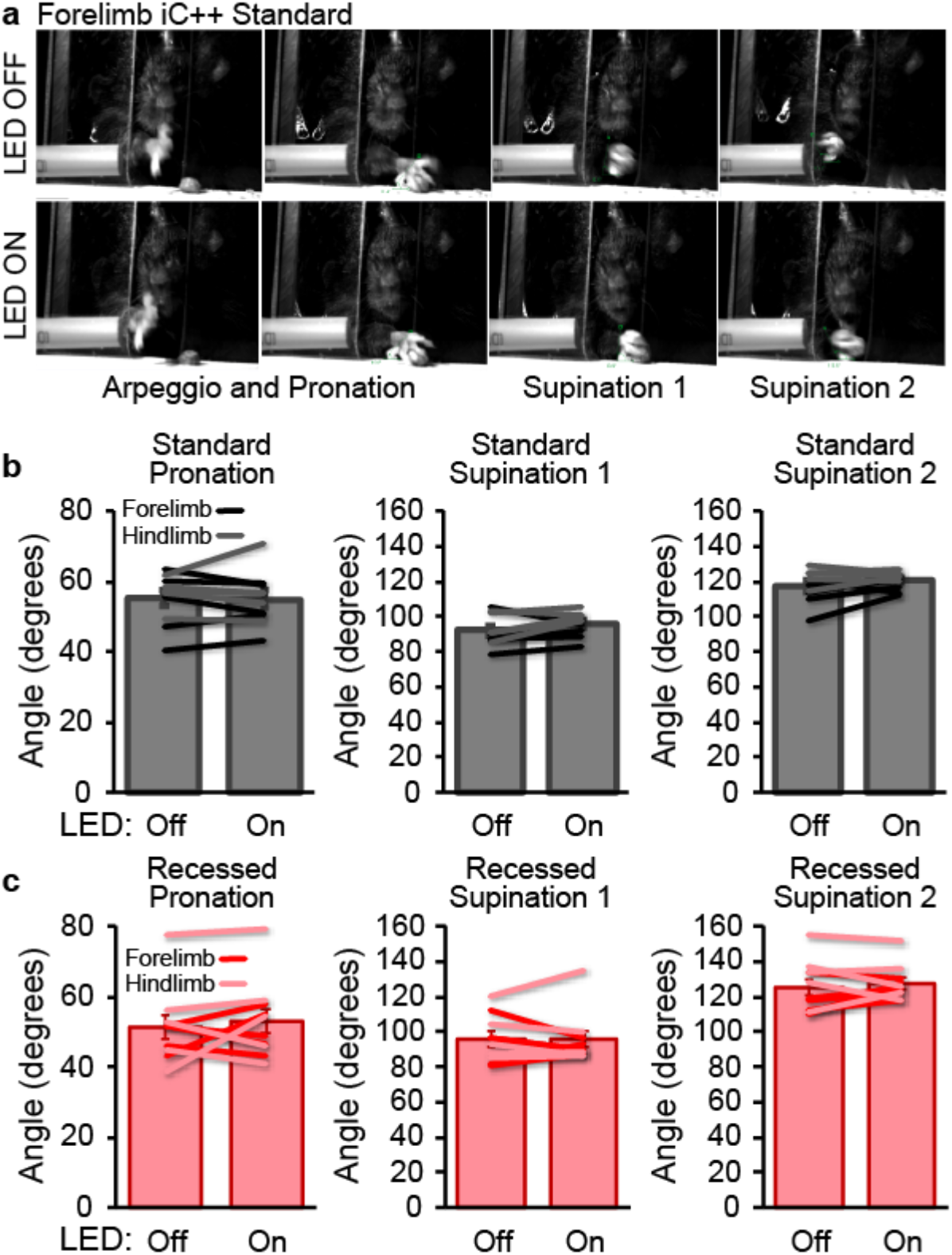
Kinematics of paw rotation during successful reach is not affected by optogenetic silencing of motor cortex with iC++. (a) Examples of pellet grasping during standard reach for a representative mouse during 473 nm LED OFF and ON sessions. iC++ silencing of neither forelimb nor hindlimb cortex alters angles of pronation, supination 1, or supination 2 during standard (b) or recessed (c) forelimb reach. Data presented as mean ± sem; *n* = 5/group.

**Supplementary Figure 3.**
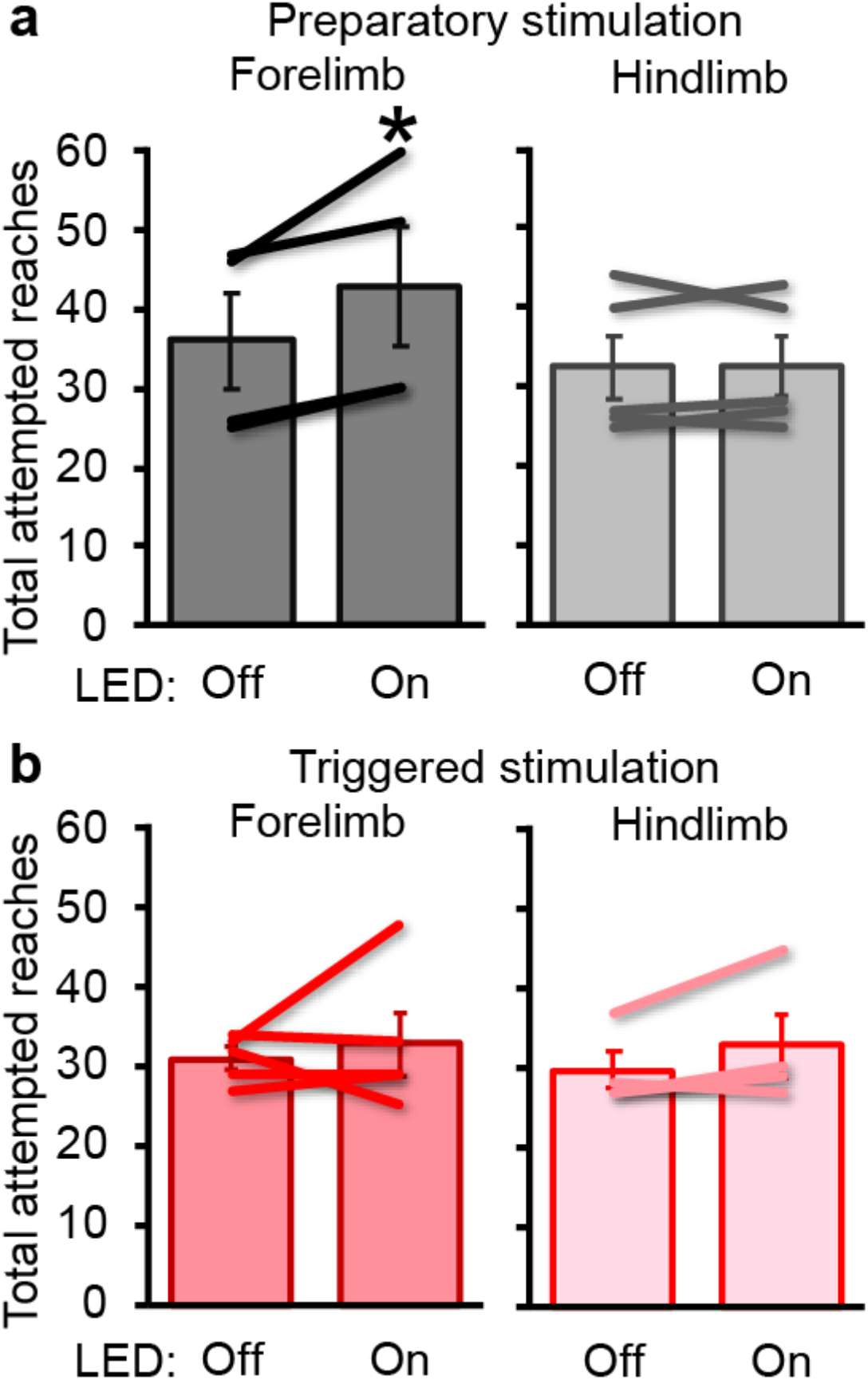
Activation of parvalbumin neurons in forelimb motor cortex impairs reach targeting when delivered prior to reach onset. The total number of attempts needed to complete 25 reaches to the pellet increased when forelimb motor cortex was silenced by ChR2-expressing parvalbumin inhibitory neurons prior to the initiation of reach, but not when silencing occurred once reach was initiated (ratio paired, two-tailed t-test *P* < 0.05). Data presented as mean ± sem; *n* = 4 (preparatory forelimb, triggered hindlimb), 5 (preparatory hindlimb, triggered forelimb).

**Supplementary Figure 4.**
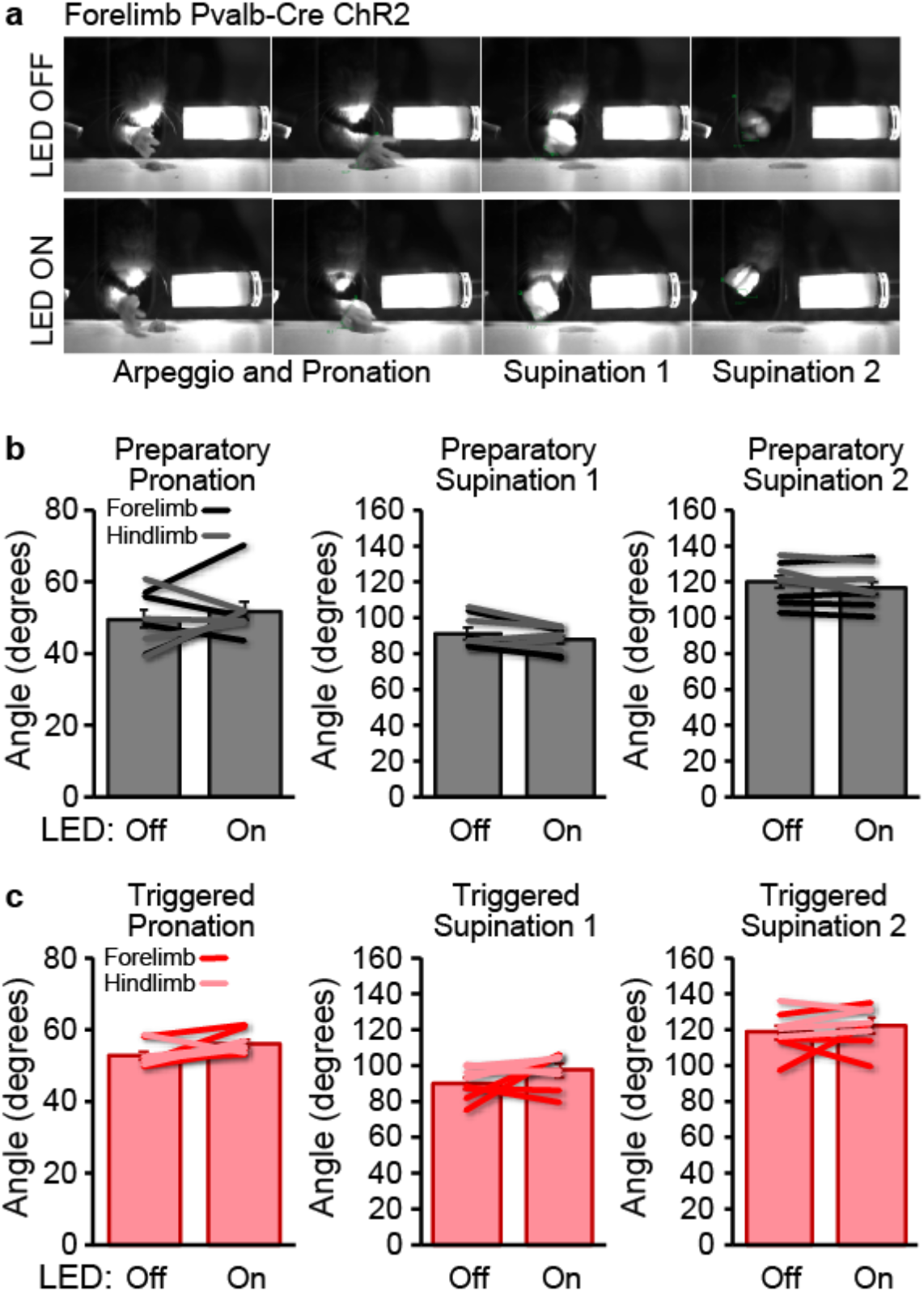
Kinematics of paw rotation during successful reach is not affected by optogenetic activation of inhibitory parvalbumin neurons in motor cortex. (a) Examples of pellet grasping during recessed reach for a representative mouse during 473 nm LED OFF and ON sessions. Parvalbumin activation had no effect on the execution of paw rotation during successful pellet retrieval. Data presented as mean ± sem; *n* = 4 (except *n* = 5 in preparatory hindlimb group).

**Supplementary Figure 5.**
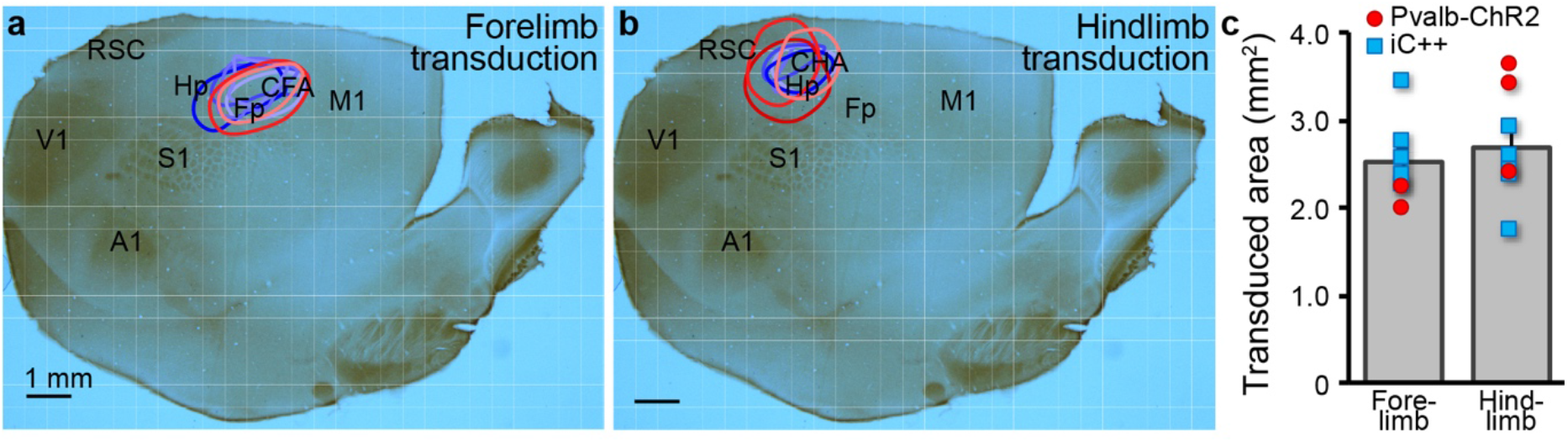
Cortical transduction of viral optogenetic constructs in mice. (a,b) Flattened mouse cortical section showing overlays of the extent of viral transduction of AAV2-Ef1a-DIO-IC++-EYFP (blue lines) and AAV2-Ef1a-DIO-hChR2(H134R)-EYFP (red lines) injections in different mice. (c) Cortical area transduced to express IC++ (blue) and ChR2 (red). Data presented as mean ± sem.

